# Engineered lignin composition in *Brachypodium distachyon* modulates the root-associated microbiome

**DOI:** 10.1101/2025.07.30.667798

**Authors:** Shweta Priya, Hsiao-Han Lin, Sannidhi Menon, Nicholas Downs, Peter F Andeer, Yang Tian, Suzanne M Kosina, Yezhang Ding, Aymerick Eudes, Trent R Northen, Aindrila Mukhopadhyay, Jenny C Mortimer

**Affiliations:** Biological Systems and Engineering Division, Lawrence Berkeley National Laboratory, Berkeley, CA, 94720, USA; Environmental Genomics and Systems Biology Division, Lawrence Berkeley National Laboratory, Berkeley, CA, 94720, USA; School of Agriculture, Food and Wine & Waite Research Institute, University of Adelaide, Adelaide, South Australia 5064, Australia

## Abstract

Lignin is a key component of the plant cell wall and a barrier to microbial interaction and biomass digestibility. While lignin biosynthesis has been engineered in grasses to reduce recalcitrance for agroindustrial applications, the consequences for root-associated microbial communities remain poorly understood. Here, we show in the model grass species *Brachypodium distachyon* that genetic modification of lignin biosynthetic genes, particularly caffeic acid *O*-methyltransferase (*COMT*), alters root lignin syringyl/guaiacyl (S/G) ratios, root exudate composition, and significantly reshapes the rhizosphere microbiome. Using a synthetic microbial community (SYNCOM) in controlled fabricated ecosystem (EcoFAB) environments, we observed a distinct reduction in *Burkholderia* colonization and enrichment of *Rhodococcus* in *COMT* mutant lines. LC-MS/MS profiling of root exudates revealed increased *p*-coumaric acid, caffeic acid and hydroxycinnamic acid amides in these lines, potentially linked to both metabolic remodeling and plant stress responses. These findings demonstrate that altering lignin composition can have profound, genotype-specific impacts on rhizosphere microbiome assembly, with implications for plant-microbe interactions, nutrient cycling, and biomass conversion strategies in bioenergy crops.

## Introduction

The plant cell wall, a dynamic and complex structure surrounding all plant cells, is a major interface between plants and their microbial environment. It comprises a matrix of polysaccharides, proteins, and, in some tissues, lignin, a phenolic polymer that contributes to structural integrity and environmental resilience. Beyond serving as a passive barrier, the root cell wall plays an active role in shaping microbial communities, influencing nutrient exchange, signaling, and symbiosis initiation [1–3]. While most studies on cell wall-microbe interactions have emphasized immunity, due to the wall’s role in pathogen defense and immune elicitation upon disruption [4,5], emerging evidence points to broader influences on the rhizosphere microbiome.

Lignin is a heterogeneous, polyphenolic compound that fortifies plant cell walls and defends against microbial invasion [5,6]. However, its recalcitrance to enzymatic degradation poses challenges for biomass conversion into animal feed and biofuels. Lignin impedes saccharification by physically blocking access to cellulose and hemicelluloses and through chemical interactions with cell wall polysaccharides [7]. Consequently, substantial efforts have been devoted to genetically modifying lignin biosynthesis to improve biomass digestibility without compromising plant growth and yield [8].

In the Poaceae (grasses), which account for ∼70% of planted crops globally, the lignin consists of guaiacyl (G) and syringyl (S) units together with a lesser amount of *p*-hydroxyphenyl (H) units with *p*-coumaric acid (CA) and ferulic acid (FA) contributing to its polymerization and complexity [7]. These lignin monomers are synthesized via the phenylpropanoid pathway, starting from phenylalanine or tyrosine and involving a series of enzymatic steps in the cytosol, before being transported outside the cell for polymerisation in the wall [9] (Figure 1). The relative abundance of syringyl to guaiacyl (S/G) monomers often correlates with biomass recalcitrance, with lower S/G ratios associated with enhanced saccharification in grasses, including the model grass *Brachypodium distachyon* [10].

**Figure 1.**
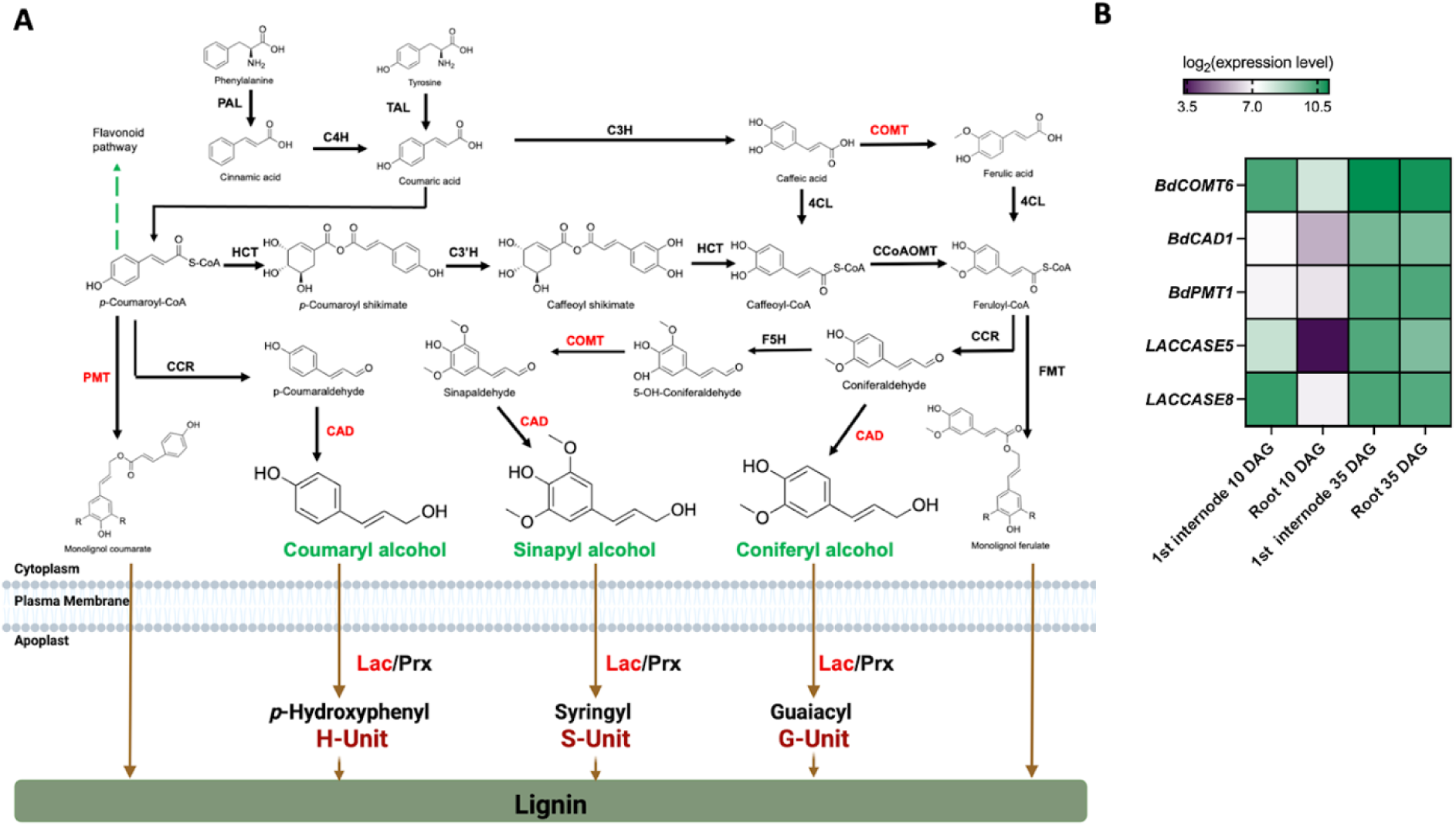
Proposed lignin biosynthetic pathway in *B. distachyon* and the expression levels of genes used in this study. (A) Phenylpropanoid pathway of lignin biosynthesis in *B. distachyon*, Phenylalanine ammonia lyase (PAL), hydroxycinnamoyl CoA:shikimate hydroxycinnamoyl transferase (HCT), *p*-coumaroyl shikimate 3′-hydroxylase (C3′H), caffeic acid/5-hydroxyferulic acid *O*-methyltransferase (COMT), 4-coumarate:CoA ligase (4-CL), caffeoyl CoA *O*-methyltransferase (CCoAOMT), cinnamoyl CoA reductase (CCR), ferulic acid 5-hydroxylase (F5H), cinnamyl alcohol dehydrogenase (CAD), laccase (Lac), peroxidase (Prx). *p*-Hydroxyphenyl (H-unit), Syringyl (S-unit), Guaiacyl (G-unit). Enzymes with altered activity in the plants used in this study are shown in red. The pathway is derived from [15] and [18], (B) Expression levels of genes for lignin biosynthesis in *B. distachyon* used in the study in 1st internode of stem and root at 10 DAG and 35DAG, DAG: days after germination. The gene expression level data were derived from [19].

Several genes at the end of the lignin biosynthetic pathway have emerged as promising targets for engineering reduced recalcitrance, including COMT (caffeic acid *O*-methyltransferase) and CAD (cinnamyl alcohol dehydrogenase) [11]. COMT is involved in conversion of caffeic acid to ferulic acid and synthesizing syringyl monolignols via methylation of 5-hydroxyconiferaldehyde [12], while CAD reduces cinnamaldehydes to their corresponding alcohols [13]. Other key enzymes include PMT (*p*-coumaroyl-CoA:monolignol transferase), which mediates *p*-coumaroylation of monolignols, and laccases (Lac), which oxidize monolignols prior to polymerization [14,15]. Disrupting the activity of these enzymes in *B. distachyon*, in particular COMT and CAD, has been previously shown to reduce lignin content, reduce the ratio of S/G and increase saccharification [16,17]. While these efforts have focused on biomass processing, relatively little is known about the broader biological consequences of such modifications, particularly at the plant-microbe interface.

Perturbations in lignin biosynthesis not only alter lignin quantity and composition but can also cause widespread metabolic reprogramming. For instance, these changes often involve the accumulation of soluble phenolics and secondary metabolites, which can influence the availability and form of the carbon source used by root-associated microbes [20–22]. In *Populus*, silencing CCR (cinnamoyl CoA reductase, see Figure 1) led to altered xylem cell wall composition and reduced colonization by endophytic *Actinobacteria* [21,23]. Similarly, changes in the shikimate pathway in transgenic switchgrass reduced root lignin and increased protocatechuate exudation, resulting in a shift in rhizosphere bacterial composition [22].

Many aromatics derived from the phenylpropanoid pathway, such as ferulic and *p*-coumaric acids, serve as carbon sources for rhizosphere microbes [24,25]. For example, flavonoids act as microbial signaling molecules, playing key roles in root colonization and the establishment of beneficial interactions with rhizobia and arbuscular mycorrhizal fungi (AMF) [26], while Tricin, a flavonoid incorporated into grass lignin, has also been shown to suppress fungal pathogens [27].

Although most studies of lignin engineering have been conducted under controlled laboratory conditions, these modifications could have unanticipated effects in more complex soil environments. For instance, altered exudate chemical composition might impact root-microbe associations critical for nutrient acquisition or stress tolerance. Understanding how cell wall engineering intersects with root microbiome assembly is therefore essential for translating these modifications into field-deployable traits.

In this study, we test the hypothesis that engineering lignin composition alters the root-associated microbiome through changes in the composition of the root cell wall and root exudates. Using a collection of *B. distachyon* lines with targeted disruptions or overexpression of lignin biosynthetic genes, we profiled lignin composition, root exudates, and microbiome structure across spatial compartments (roots, rhizosphere, and bulk sand) using a defined synthetic bacterial community (SYNCOM). This approach allows us to isolate the effects of plant genotype on microbial community assembly and offers insights into the metabolic and ecological consequences of lignin modification.

## Materials and Methods

### Plants lines and growth in EcoFABs

*Brachypodium distachyon* accession Bd21-3 (wild type, WT) and engineered lines with modified lignin biosynthesis used in this study are shown Table 1. Seeds were stored at room temperature until use. For use, seeds were dehusked, and surface sterilized in 70% (v/v) ethanol for 2 min, followed by 10% (v/v) sodium hypochlorite for 5 min, and rinsed five times with sterile ultrapure (18.2 MΩ) water. Plants were grown as previously described [28]. Sterilized seeds were placed on 0.5× Murashige and Skoog (MS) basal medium with vitamins and 1.0 g/L MES (PhytoTech Labs, Lenexa, KS, USA) without sucrose solidified with 1.5% (w/v) Phytoagar and stratified at 4 °C in the dark for 2 days. Plates were incubated in a growth chamber at 24 °C with 60% relative humidity and a 16 h light/8 h dark photoperiod (light intensity: 120 μmol m^-2^s^-1^). After 3-4 days, seedlings were transferred to EcoFAB 2.0 chambers [29] using sterile forceps. EcoFABs were filled with 10 g sterile quartz sand (50-70 mesh; Sigma-Aldrich, St. Louis, MO, USA) and 7 mL of 0.5× MS liquid medium without sucrose. Plants were irrigated with sterile water every 2-3 days and harvested at 3 weeks post-germination.

**Table 1.** Brachypodium distachyon WT and engineered lines used in the study.

| *B. distachyon* line | Genotype | References |
| --- | --- | --- |
| Bd21-3(WT) | Wild type | [30] |
| COMT5139 | Mutation in *BdCOMT6* (*Bradi3g16530*) gene in *Bd5139* | [16] |
| CAD4179 | Mutation in *BdCAD1* (*Bradi3g06480*) gene in *Bd4179* | [17] |
| CAD7591 | Mutation in *BdCAD1* (*Bradi3g06480*) gene in *Bd7591* | [17] |
| PMT6063 | Mutation in *Bdpmt-1* (*Bradi2g36910*) gene | [14,17] |
| *lac5lac8* | Mutations in *LACCASE5* (*Bradi1g66720*) and *LACCASE8* (*Bradi2g23370*) genes (*Bd4442/Bd5731*) | [31] |
| *PMT OX* | Overexpression of *Bdpmt-1* (*Bradi2g36910*) gene under the control of the CaMV 35S promoter. | [14] |

### Root Lignin Composition Analysis

Roots were separated, flash-frozen in liquid nitrogen, lyophilized, and ground to a fine powder. Lignin monomer composition was assessed using pyrolysis gas chromatography mass spectrometry (Py-GC/MS) as previously described [32]. Powdered samples (0.5 mg) were pyrolyzed at 650 °C using the pyroprobe 5200 (CDS Analytical, Inc., Oxford, PA, USA) connected to a gas chromatography mass spectrometry (GC/MS) system (Agilent 6890) composed of a Trace GC Ultra and a Polaris-Q MS (Thermo Electron Corporation, Waltham, MA, USA) equipped with a TR-SMS column (60 m 0.25 mm ID 0.25 lm) and operated in split mode (40 mL min^−1^) using He gas as carrier. The chromatograph program was set as follows: 5 min at 50 °C, followed by an increase of 5 °C min^−1^ to 300 °C, finally maintained at 300 °C for 5 min. Lignin-derived fragments were identified via the NIST08 library. Relative abundances of syringyl (S), guaiacyl (G), and *p*-hydroxyphenyl (H) units were compared from peak areas. S/G ratios were calculated by dividing total S peak area by total G peak area.

### Root Exudate Analysis

For LC-MS/MS analysis of root exudates, the surface sterilized seeds were germinated in the same conditions as above. The Magenta vessels (Fisher Scientific, NH, US) were rinsed 5 times with ultrapure water and sterilized by autoclaving. Two *B. distachyon* seedlings (2 days old) were placed onto a sterilized floating holder and then transferred to each vessel pre-filled with 40 mL of 0.5× MS medium (same as used before) in the growth chambers under the same conditions as described above. Five biological replicates of each plant line (WT, COMT5139 and *PMT OX*) were used for the root exudates collection. Three-week-old plants were then carefully removed from the vessels together with the floating holder and 15 mL 0.5× MS medium (with dissolved root exudates) were collected in 50 mL conical tubes and stored at −80°C for further analysis.

### Metabolomics analysis

Frozen root exudates were freeze dried and then suspended in 1 mL of methanol. Tubes were vortexed twice for 10 sec, then bath sonicated in ice water for 15 min. Solutions were transferred to 2 mL snap cap tubes, then centrifuged at 10,000 rcf for 5 min at 10 ℃. Supernatants were collected into a new tube, dried by vacuum concentration and stored at −80 ℃. On the day of LC-MS analysis, dried frozen extracts were resuspended in 150 µL of methanol with internal standard mix (Table S2). These were vortexed, sonicated and centrifuged as before, however, the final supernatant was then filtered through 0.22 µM microcentrifugal filters. Filtrates were transferred to amber glass LC-MS/MS vials and capped immediately. Metabolites were separated and eluted using reverse phase chromatography and detected using a Thermo Orbitrap Exploris 120 Mass Spectrometer (LC-MS/MS parameters are available in Table S2). Raw data files are available at https://gnps2.org/status?task=446d069a99fe49cdb11fe5ddf66bd800. Targeted metabolomics analysis was performed using Metatlas (https://github.com/biorack/metatlas) [33]; briefly, m/z, retention time and fragmentation spectra from samples were compared with pure reference standards for identification of detectable metabolites (Table S3). Untargeted features and MS2 spectra were extracted using MZmine 3.7.2 [34], and then analyzed on Global Natural Product Social Molecular Networking (GNPS) site using the feature based molecular networking workflow [35]. GNPS outputs are available at https://gnps2.org/status?task=74996ed8d32c42f693d3905e9e518b06 (negative mode) and https://gnps2.org/status?task=ac1e0917315546d09f70a728f0ebfe2e (positive mode). Putative annotations (Table S4) were determined by filtering untargeted features to include only those with GNPS spectral matches with an MQscore >0.7, ppm <0.7, an adduct match to the correct sample polarity, retention time between 1 and 11 minutes, at least one sample with an intensity 100,000 counts greater than the maximum intensity in the controls; additionally lipids (not optimized on this method) and contaminants were removed from the final list. Peak heights (Table S5) of both targeted metabolite identifications and putative annotations from GNPS matching were compared between WT and lignin engineered lines using one way ANOVA followed by Dunnett’s multiple comparison test. The Principal Component Analysis (PCA) plot was created using the ‘matplotlib’ library in python [36]. A contingency plot was generated using the log2 transformed fold changes in peak heights of engineered lines relative to the WT. GraphPad Prism 10 (GraphPad Software, San Diego, CA, USA) was used to make the contingency plot and perform statistical analysis.

### Synthetic community (SYNCOM) preparation and inoculation

The bacterial strains used in this study (Table 2) were obtained from a *Panicum virgatum* (switchgrass) field in Oklahoma, United States [37]. Strains were cultured at 30 °C at 180 rpm using Reasoner’s 2A (R2A) medium [38,39], 10 % (v/v) strength R2A (0.1× R2A), Luria-Bertani medium (Thermo Fisher Scientific), and yeast-malt extract medium as indicated in Table 2. To test their ability to grow on the plant medium, bacterial strains were streaked simultaneously on 0.1× R2A agar and 0.5× MS agar. Bacterial strains (except *Bradyrhizobium* OAE829) were grown for 48 h in 10 mL of culture medium as described earlier. *Bradyrhizobium* OAE829, which does not grow well in liquid medium, was grown on several 0.1× R2A agar plates for 10 days, and the bacterial mass was collected using sterile loops, pooled, and resuspended in 0.5× MS. Cultures were centrifuged at 4,000 × g for 20 min and washed twice with 0.5× MS. Then the OD_600_ of each culture was measured and the 17 strains were pooled such that each had a final OD_600_ equal to 1. This SYNCOM liquid mixture was diluted 100 times, and this served as the inoculant. Inoculant (70 µL) was applied directly to each EcoFAB root growth chamber which had been prefilled with 7 mL of 0.5× MS medium (Figure 2).

**Figure 2.**
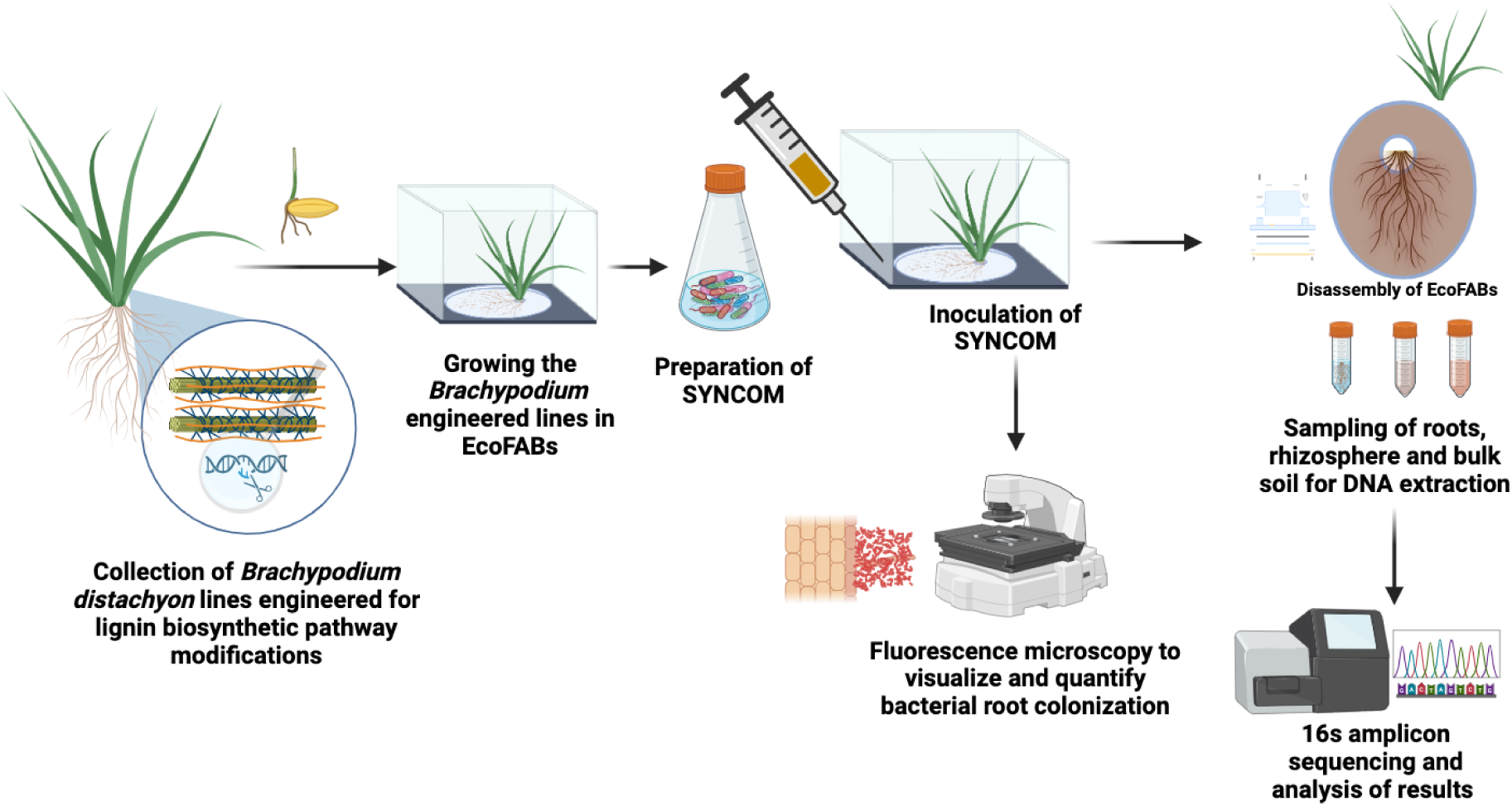
Experimental workflow for examining the impacts of lignin modifications in *B. distachyon* on microbial community (SYNCOM) composition and assembly in the roots, rhizosphere and bulk sand. The figure was created with biorender.com.

**Table 2.** List of bacterial strains in the SYNCOM.

| SYNCOM Bacteria name | Medium used to cultivate |
| --- | --- |
| *Arthrobacter* OAP107 | R2A |
| *Bacillus* OAE603 | TSB |
| *Bosea* OAE506 | R2A |
| *Bradyrhizobium* OAE829 | 0.1× R2A |
| *Brevibacillus* OAP136 | R2A |
| *Burkholderia* OAS925 (renamed *Paraburkholderia graminis* OAS925) | R2A |
| *Chitinophaga* OAE865 | R2A |
| *Lysobacter* OAE881 | R2A |
| *Marmoricola* OAE513 | R2A |
| *Methylobacterium* OAE515 | LB |
| *Mucilaginibacter* OAE612 | R2A |
| *Mycobacterium* OAE908 | R2A |
| *Niastella* OAS944 | R2A |
| *Paenibacillus* OAE614 | R2A |
| *Rhizobium* OAE497 | R2A |
| *Rhodococcus* OAS809 | R2A |
| *Variovorax* OAS795 | R2A |

### Root Microbiome Sample Harvesting

Plants were harvested 3 weeks after inoculation. Shoots were excised, and fresh weight recorded. Roots were gently tapped to remove loosely attached sand, then transferred to a 50 mL conical tube containing 5 mL 1× PBS. Tubes were vortexed twice (10 s each) to detach rhizosphere sand. Separated sand was collected into separate conical tubes for bulk analysis. Roots were then surface sterilized using 3% (v/v) H_2_O_2_ for 30 sec, followed by washing with 100% ethanol for 30 sec and then with 6.15% (v/v) NaOCl with 2-3 drops of Tween-20 per 100 mL for 3 min, followed by a final wash with 3% (v/v) H_2_O_2_ for 30 sec. Roots were then transferred to 50 mL conical tubes containing 25 mL sterile water. Surface sterility was confirmed for all samples by touching a subsampled root from each collection onto LB plates and incubating overnight at 30 °C. Roots were then transferred into a pre-weighed 2 mL tube containing 0.64 cm steel balls, re-weighed and stored at −20 ℃ for further analysis.

### 16S rRNA gene DNA extraction and sequencing

Bacterial DNA was extracted using the DNeasy PowerSoil HTP 96 Kit (Qiagen, Redwood City, CA, USA) according to the manufacturer’s instructions. Bulk sand and rhizosphere samples were prepared by adding 250 mg of the collected substrate to a PowerBead plate (Qiagen) and homogenizing with a TissueLyser II (Qiagen) at a frequency of 20 Hz for 10 min (twice). Root samples were homogenized with a 0.64 cm precision ball using the TissueLyser II (Qiagen) at 30 Hz for 3 min (twice). Homogenized root samples were transferred to a PowerBead plate, and DNA was extracted with the other samples. To extract DNA from pure SYNCOM culture (T0 inoculum), the SYNCOM pellet was added into a PowerBead Pro Tube.

DNA concentrations were quantified by a Quant-iT dsDNA high-sensitivity kit (Thermo Fisher Scientific, Waltham, MA, USA) and normalized to 0.3 ng/µL. DNA template was added to a polymerase chain reaction to amplify the V4/V5 16S gene region using 515F/926R primers based on the Earth Microbiome Project primers (<u>Parada et al. 2016</u>; <u>Quince et al. 2011</u>) but with in-line dual Illumina indexes (<u>Price et al. 2018</u>; <u>Sharpless et al. 2022</u>). The amplicons were sequenced on a MiSeq with 600-bp MiSeq Reagent Kit v3 (Illumina, San Diego, CA, USA).

### Quantification of rhizosphere colonization by *Burkholderia* OAS925 and fluorescent microscopy

To determine the differential rhizosphere colonization by *Burkholderia* OAS925 among the SYNCOM bacteria, we inoculated *Brachypodium* grown with 16 SYNCOM bacteria (as listed in Table 2 except the *Burkholderia* OAS925) and OAS925 harboring the plasmid pJ23100-RFP [40] with genes for kanamycin resistance and red fluorescent protein (RFP). Plants were harvested and rhizosphere samples were collected after 2 weeks post inoculation following the same method described above. One mL of the rhizosphere aliquot (collected after the sand settled in the 1×PBS) was serially diluted 100× and plated on R2A media plates supplemented with 10 µg/mL of kanamycin to isolate and select the *Burkholderia* OAS925 from the SYNCOM. The colonies were counted in 48 h as CFU per mL per g of root. GraphPad Prism 10 (GraphPad Software, San Diego, CA, USA) was used to perform statistical analysis using one-way ANOVA and to make the plot. The images of the inoculated roots were taken with an EVOS M7000 fluorescence microscope (Thermo Fisher Scientific, Waltham, MA, USA) using objectives with 2X to 40X magnification.

### 16s rRNA amplicon sequencing data analysis and visualization

The 16S rRNA gene amplicons were analyzed as described in [28]. Beta diversity was calculated by Bray-Curtis distance using R 4.2.2 with package ‘vegan 2.6.4’, which was used to create the Principal coordinates analysis (PCoA) plots using the ‘ggplot2’ package [41]. Permutational multivariate ANOVA (PERMANOVA) was calculated using ‘adonis2’ in the ‘vegan 2.6.4’ package. Pairwise comparisons were performed using DESeq2 v1.38.3 [42].

## Results

### *B. distachyon* lignin mutants with altered aerial lignin composition also have altered root lignin composition

In this study, we utilized a collection of previously published *B. distachyon* lines with reported altered stem lignin composition (Table 1). Further analysis of gene expression revealed that the majority of lignin-related genes upregulated in the stem are also upregulated in maturing root tissues (Figure 1B, Table S6). To examine if these lines also exhibit altered root lignin composition, we performed pyrolysis-GC/MS (Py-GC/MS) analysis on root tissues harvested from 21-day-old plants. Wild-type (WT, Bd21-3) roots exhibited a syringyl-to-guaiacyl (S/G) monomer ratio of 0.85 ± 0.04, consistent with the previously reported ranges for grasses [43,44].

All lignin-engineered lines exhibited altered root S/G ratios compared to WT (Table 3). COMT5139 showed the greatest reduction in S/G (0.40 ± 0.03; *p* < 0.005), followed by PMT6063 (0.50 ± 0.03; *p* < 0.05) and CAD4179 (0.61 ± 0.07). In contrast, the overexpression line, PMT-OX, exhibited a slightly elevated S/G ratio (0.98 ± 0.08), ∼15% above WT. These results demonstrate that the targeted genes in lignin-engineered lines are responsible for lignin biosynthesis in both above-ground and root tissues, leading to modifications of lignin composition in the roots of these lines.

**Table 3.**
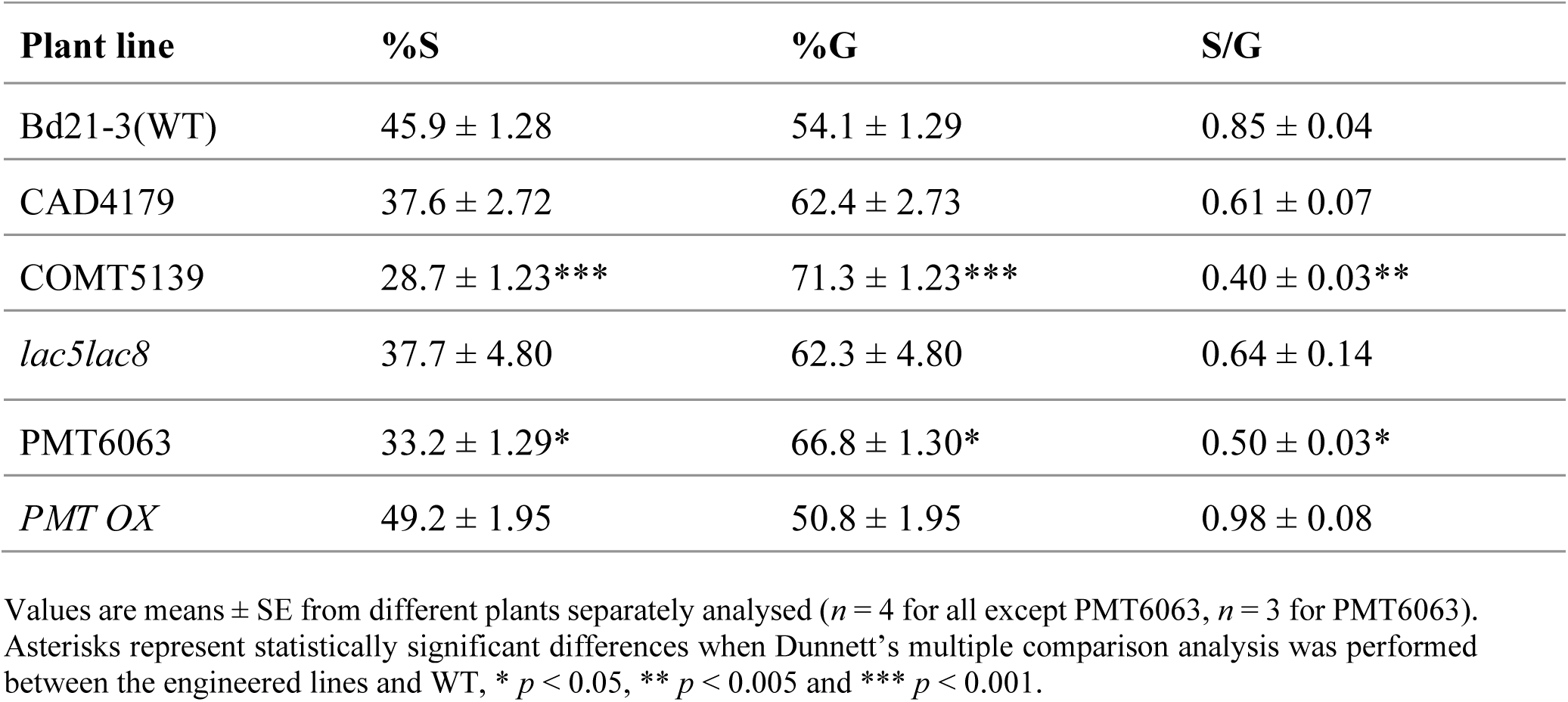
Root lignin monomer composition in all the *Brachypodium* lines used in the study.

### Microbiome composition is shaped by both plant genotype and spatial niche

To examine the impact of lignin modification on the root-associated microbiome, we selected three genotypes representing a spectrum of root S/G ratios: COMT5139 (lowest S/G), PMT6063 (low S/G), and *PMT OX* (high S/G). Plants were grown in EcoFABs and inoculated with the SYNCOM, and microbial DNA was extracted from root, rhizosphere, and bulk sand compartments after three weeks for 16S rRNA amplicon sequencing (Figure 2).

Principal coordinates analysis (PCoA) of Bray-Curtis dissimilarities revealed clear separation by both plant genotype and sample type (Figure 3). The first two axes accounted for 68.76% of the total variance (PCoA1: 56.32%; PCoA2: 12.43%). PERMANOVA confirmed significant effects of genotype (*p* = 0.004) and sample type (*p* < 0.0001) on microbiome beta diversity. Root samples clustered separately from all other samples, suggesting niche-specific assembly patterns. However, it should be noted that read counts for the root samples were low (between 70 - 3000) and so would need further investigation with larger sample sizes to draw robust conclusions.

**Figure 3.**
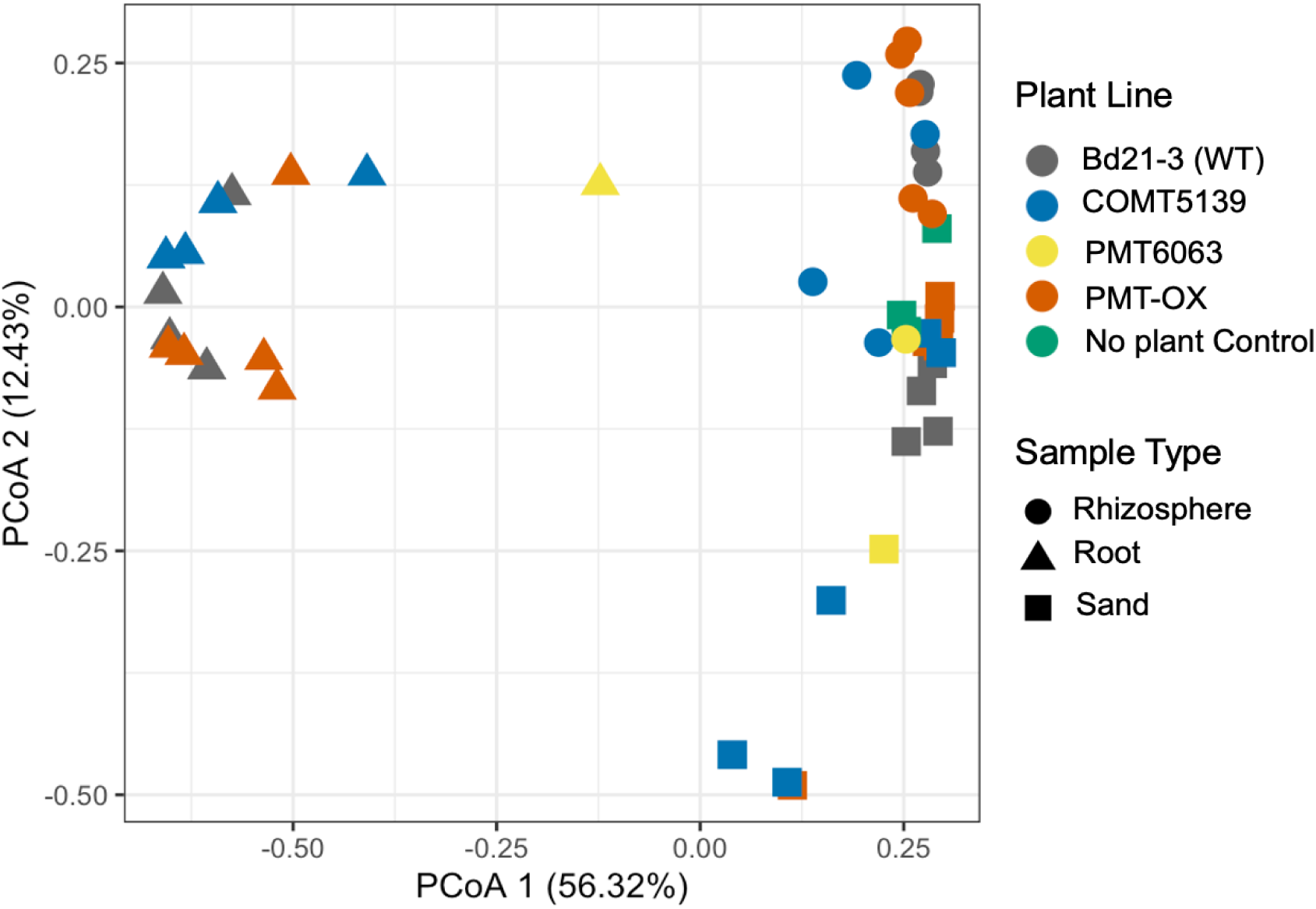
Principle coordinate analysis (PCoA) plot of microbiome beta diversity across all sample types and all plant genotypes. Samples are colored by the plant line and shape reflects the sample type.

### The rhizosphere microbiome is most strongly affected by lignin modification

We next tested whether microbiome composition varied by genotype within each spatial compartment. PERMANOVA pairwise comparisons revealed a significant genotype effect in the rhizosphere (*p* = 0.0102), but not in the bulk sand (*p* = 0.065) or root samples (*p* = 0.13) (Table 4). As noted above, the lack of significance in root samples is likely due to low sequencing depth. Among the genotypes tested, COMT5139 exhibited the most significant shift in rhizosphere community composition relative to WT (PERMANOVA: *p* < 0.05), while *PMT*-*OX* showed no significant difference (Table S1).

**Table 4.** PERMANOVA pairwise comparisons between sample types and plant genotypes. Statistically significant (*p* < 0.05) values are shown in bold format with asterisks (*).

| PERMANOVA comparisons with plant genotype | R<sup>2</sup> | *p*-value |
| --- | --- | --- |
| All sample types | 0.07278 | 0.06739 |
| Sand | 0.31029 | 0.06529 |
| Rhizosphere | 0.39882 | **0.0102\*** |
| Root | 0.26067 | 0.1298 |

To further validate these results, we carried out a larger scale experiment where we analyzed only the rhizosphere microbiome, but included additional lines with altered lignin (CAD4179, *lac5lac8* and CAD7591), in addition to the original mutant lines (COMT5139 and PMT6063). Again, only COMT5139 showed a significant rhizosphere shift versus WT (*p* < 0.05), confirming that in our experimental system, the root compositional changes that arise as a result of the *COMT* mutation specifically drive reproducible changes in microbial assembly (Figure 4, Table S1).

**Figure 4.**
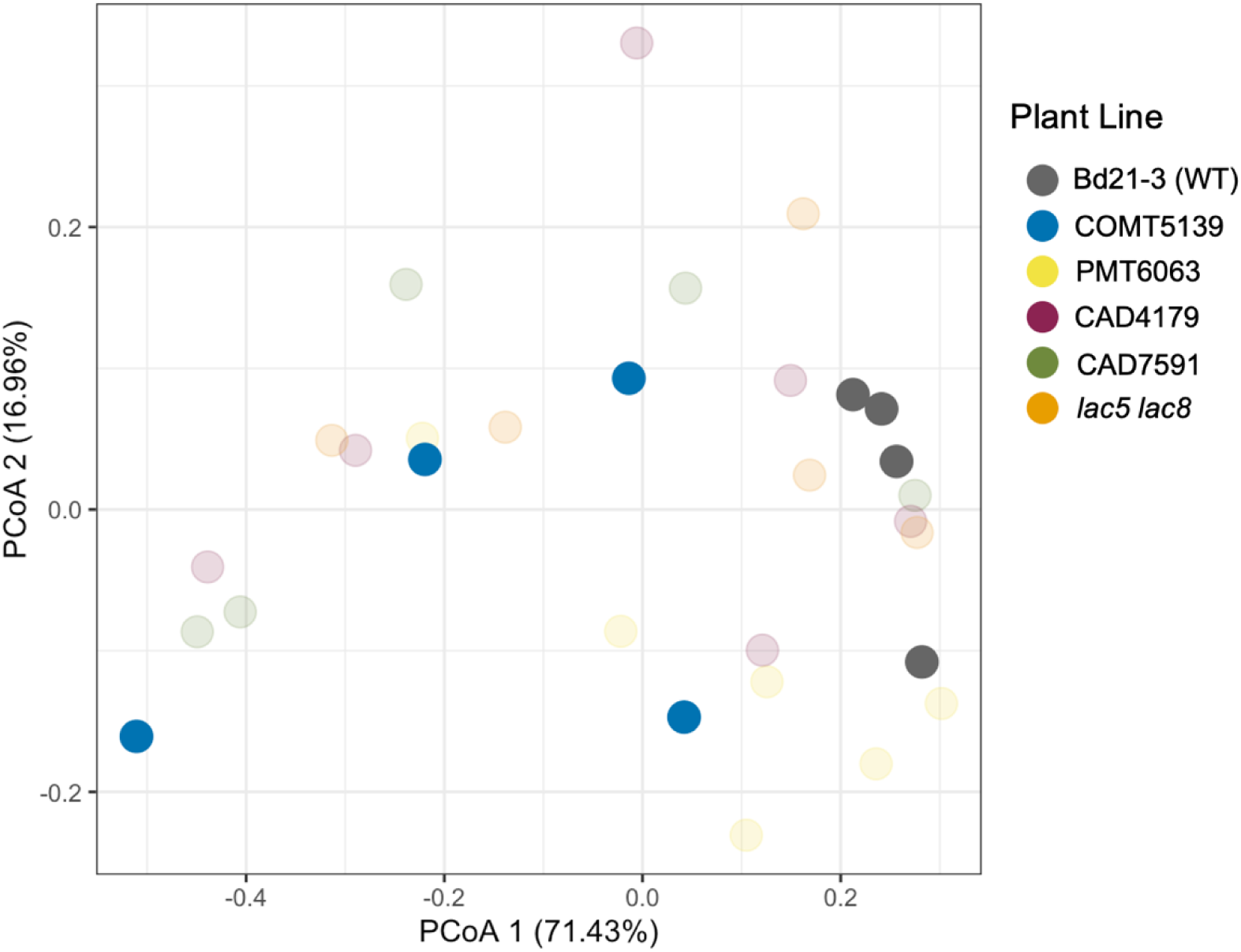
Principle coordinate analysis (PCoA) of the rhizosphere microbiome beta diversity across all the plant lines. Samples are colored by plant lines and shaped by plant type (WT or transgenic). The solid circles show the plant lines that had a significantly different rhizosphere microbiome diversity than WT (PERMANOVA pairwise comparison with WT, *p* < 0.05) and transparent circles are the lines that showed no significant differences in rhizosphere microbiome diversity than WT (PERMANOVA pairwise comparison with WT, *p* > 0.05).

### Specific members of the SYNCOM are differentially abundant in the rhizosphere of COMT5139 compared to WT

To identify which microbial taxa were responsible for the observed differences in community composition, we performed DESeq2 differential abundance analysis on rhizosphere samples from WT and COMT5139 plants. Two SYNCOM members were significantly differentially abundant: *Burkholderia* OAS925 was significantly depleted in COMT5139 rhizospheres (*Padj* = 0.0356), while *Rhodococcus* OAE809 was significantly enriched (*Padj* = 0.0356; Figure 5A). The significantly lower colonization of *Burkholderia* OAS925 in COMT5139 was further validated by inoculating the seedlings with the SYNCOM along with a strain of OAS925 expressing RFP (and kanamycin resistance for selective isolation) and plating the resulting rhizosphere microbial community on selective media plates. The results showed that the colonization of OAS925 was 3× lower in the rhizosphere of COMT5139 compared to that of WT, however, no significant difference in OAS925 colonization was observed for the *PMT OX* rhizosphere (Figure 5B). To further explore this, we used an RFP-expressing line of OAS925 inoculated as part of the SYNCOM to look at the spatial distribution of the microbes across the root surface in EcoFAB-grown seedlings. In agreement with the CFU data, we observed further decreased root colonization of OAS925 in COMT5139, while the *PMT OX* did not show very substantial difference in root colonization compared to the WT (Figure 5C).

**Figure 5.**
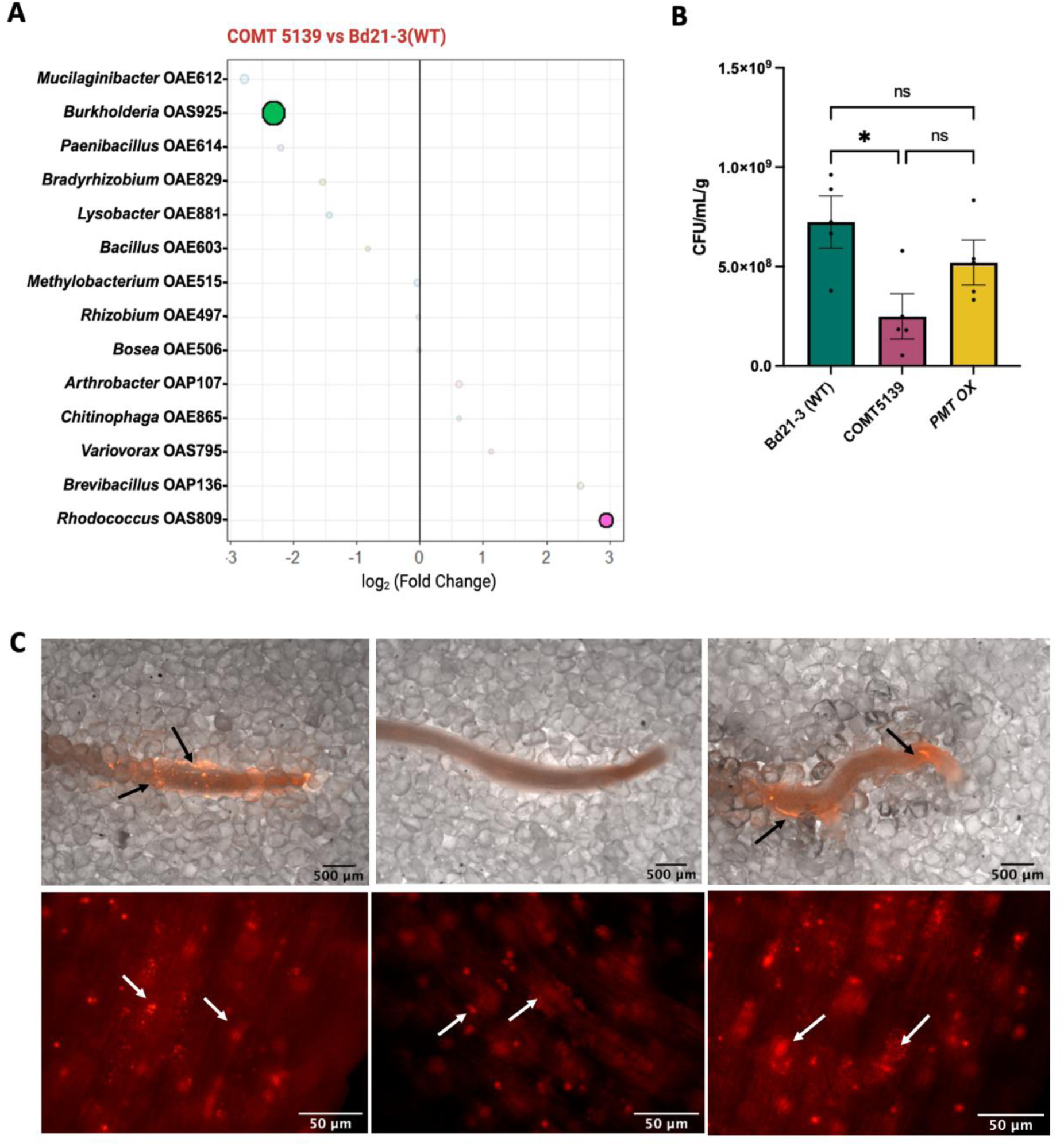
Differential abundance and colonization of members of SYNCOM in WT vs COMT5139. (A) Pairwise comparison (DESeq2 analysis) to reveal the differentially abundant bacteria in the rhizosphere of COMT5139 than WT. The *x*-axis represents the log2 fold change and the positive values indicate higher abundance in the rhizosphere, significantly differentially abundant bacteria (*Padj* < 0.05) are shown in solid colors. Circle size represents the base mean. Statistical analysis was performed using DESeq2 by a Wald test and corrected using the Benjamini-Hochberg method for multiple testing. (B) Rhizosphere colonization by *Burkholderia* OAS925 in WT, COMT5139 and *PMT OX*. Statistical analysis was performed using one-way ANOVA, * represents significance at *p* < 0.05 and ns represents non-significant differences. (C) Fluorescent microscopy showing the root colonization by RFP-expressing OAS925 in WT (first column), COMT5139 (middle column) and *PMT OX* (third column) when inoculated with other SYNCOM members, arrows pointing to OAS925 (red) present on the root surface and in the root tissues as an endophyte (white arrows).

### Loss of COMT activity alters root exudate composition

To explore whether changes in observed rhizosphere microbial composition correlated with differences in root exudation, we profiled root exudates from WT, COMT5139, and *PMT OX* lines using LC-MS (Figure 6). Principal component analysis (PCA) of all detected metabolites revealed that COMT5139 exudate profiles clustered separately from both WT and *PMT OX*, which were more similar to each other. The first two principal components accounted for approximately 82% of the total variance. Negative controls (no-plant samples) formed a distinct group, confirming that metabolite profiles reflected plant-derived compounds.

**Figure. 6.**
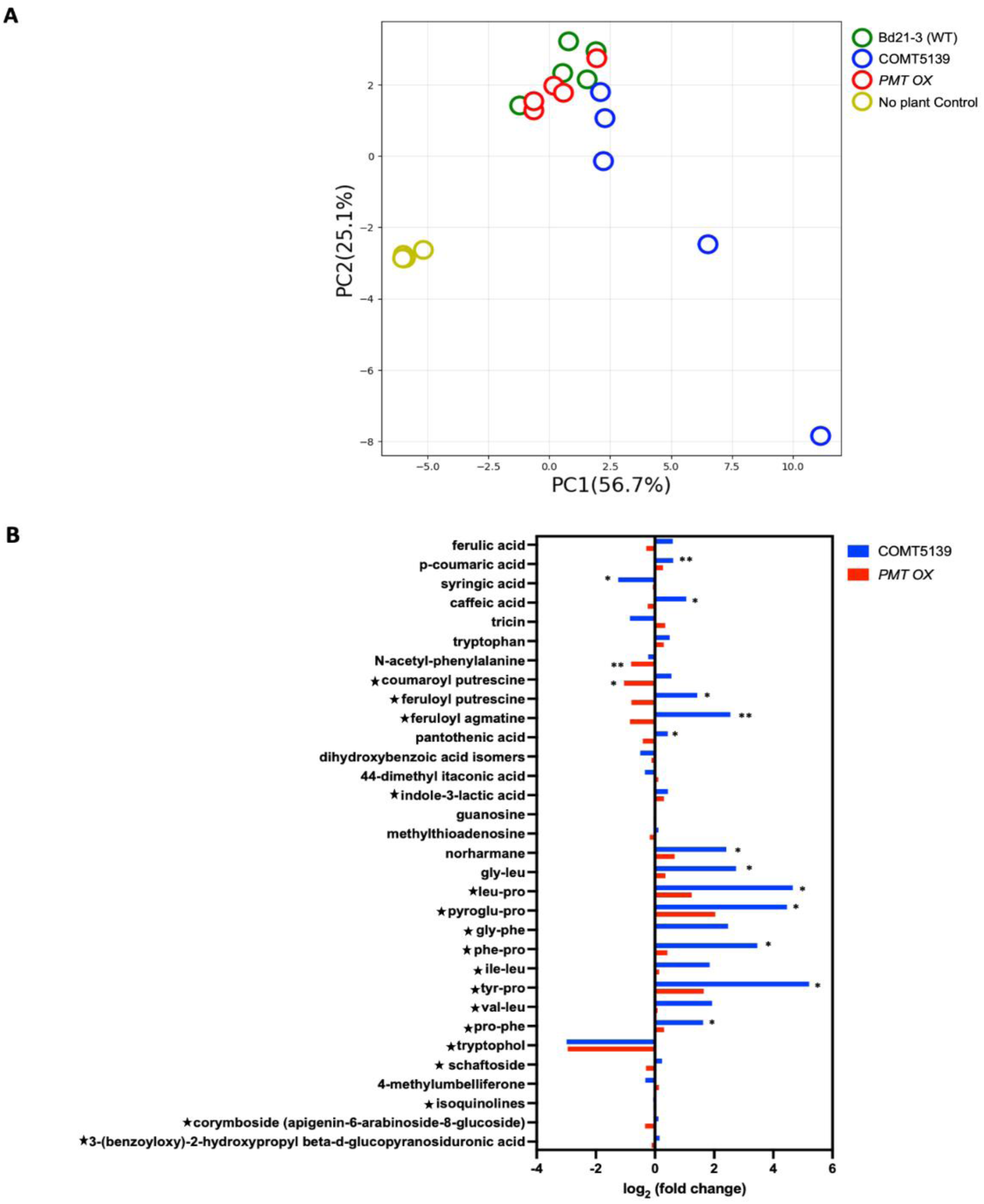
Root exudate compositional analysis of WT, COMT5139 and *PMT OX*. (A) Principal Component Analysis (PCA) of root exudate compositions of all the plant lines. (B) Contingency plot showing the differences of root exudate composition of COMT and *PMT OX* compared to WT. The star (★) in front of the compounds denotes the putatively annotated compounds. Dunnett’s multiple comparison test was performed to find the significant differences between WT and the lignin engineered lines (COMT5139 and *PMT OX*) for individual root exudate compound (n = 5), \**p* < 0.05, \*\**p* < 0.005 (B).

Several metabolites were substantially altered in COMT5139 exudates compared to WT. These included an increase in the phenolic acids *p*-coumaric acid, caffeic acid and feruloyl putrescine, as well as hydroxycinnamic acid derivatives such as feruloyl agmatine. Elevated levels of norharmane and multiple dipeptides were also observed. Conversely, the abundance of syringic acid, a downstream product of S-type monolignols, was significantly decreased in COMT5139. A substantial but insignificant (p > 0.05) decrease in the levels of tricin was also observed in COMT5139 root exudates. In contrast, the exudate profile of *PMT OX* showed only modest differences from WT, including increased levels of certain terpene lactones and reduced *N*-acetyl-phenylalanine and feruloyl putrescine. Together, these results indicate that the *COMT* mutation induces major metabolic reprogramming in the root, including alterations in both phenolic metabolism and amino acid derivatives, which may contribute to the observed shifts in rhizosphere microbiota.

## Discussion

This study revealed that *B. distachyon* lines engineered primarily for stem lignin exhibited significant alterations in root lignin composition. Additionally, similar gene expression levels in stem internode and mature roots confirm that the same set of genes are involved in lignification in both tissues (Figure 1B, Table S6). Given that roots are pivotal for microbiome assembly and symbiosis, comprehending the impact of lignin engineering on the microbiome composition underground is essential. To explore this, we combined the use of synthetic microbial community (SYNCOM) with a fabricated ecosystem (EcoFAB) to simplify the complexities of the plant-soil-microbiome nexus, and to test the effects of these defined compositional changes to the root lignin biosynthesis pathway.

### Altered lignin composition influences rhizosphere microbiome assembly

Our results demonstrate that genetic modification of lignin biosynthesis in *B. distachyon* can markedly alter the composition of the rhizosphere microbiome. Specifically, changes in the S/G monolignol ratio were associated with distinct microbial communities in the rhizosphere but not in the root endosphere or surrounding bulk medium. This genotype-specific microbiome effect supports previous findings that plant genetics influence microbial recruitment, particularly in root-associated niches [45], [46], [47]. However, few studies to date have explicitly linked cell wall engineering to microbiome shifts, particularly in grasses [22,48].

Our observed significant effect on the rhizosphere microbiome, but not that of the root interior, contrasts with studies in *Populus* where CCR silencing impacted the root endophytic microbiome with minimal effect on rhizosphere composition [21]. Conversely, studies in switchgrass with modified lignin content have reported broader changes in the microbiome of both roots and rhizosphere [22]. These inconsistencies may reflect species-specific cell wall structures, exudate profiles, or differences resulting from experimental configurations/systems.

The changes in lignin composition, particularly reductions in S-type monolignols, could alter the biochemical environment around roots, either by changing cell wall permeability, exudate composition, or carbon allocation. These changes, in turn, likely influence the niche preferences and colonization success of specific microbes in the rhizosphere.

### The COMT mutation consistently reshapes microbial composition via metabolic reprogramming

Among the engineered plant lines, COMT5139 displayed the most consistent and pronounced impact on rhizosphere microbiome composition. Notably, COMT5139 represents one of the earliest disruptions in the lignin biosynthesis pathway among our collection of lignin mutants (Figure 1). COMT is a multifunctional enzyme primarily responsible for methylation of caffeic acid and 5-hydroxyconiferaldehyde to produce ferulic acid and S-type monolignols respectively. In line with previous studies on stem tissue [16], our Py-GC/MS data confirm a substantial reduction in S/G ratio in COMT5139 roots, indicating that genetic perturbations in lignin biosynthesis manifest throughout the plant. Moreover, we also observed that the *COMT* gene is equally expressed in both stem and mature roots (Figure 1B).

Accompanying this structural change in lignin, we observed extensive metabolic reprogramming in root exudates. COMT5139 plants exuded significantly more *p*-coumaric acid, caffeic acid and hydroxycinnamic acid amides, alongside elevated levels of dipeptides and stress-related metabolites. These shifts in exudate are consistent with the idea that reduced lignin synthesis may redirect phenylpropanoid flux toward soluble intermediates, leading to altered exudate composition [20,49].

Such metabolic changes could affect microbial recruitment through multiple mechanisms, including differential carbon source availability, altered signaling molecules, or the induction of plant immune responses. Our results show that even in a simplified system using a synthetic community, these changes are sufficient to reproducibly shift microbial composition in the rhizosphere.

### COMT5139 drives consistent shifts in specific microbial taxa

Two taxa in the synthetic community were consistently and significantly altered in the rhizosphere of COMT5139 plants. *Burkholderia* OAS925 was significantly depleted, while *Rhodococcus* OAE809 was enriched, suggesting that lignin composition or its metabolic consequences influence microbial competitive dynamics.

This pattern aligns with known substrate preferences. *Burkholderia* species are frequently enriched in lignin-rich environments and are known to catabolize S-type lignin derivatives such as syringaldehyde and syringic acid [50,51]. In our study, the reduced abundance of syringic acid in COMT5139 exudates may have led to diminished colonization by *Burkholderia*. In contrast, *Rhodococcus* species are more commonly associated with G-type lignin derivatives and *p*-coumaric acid catabolism [52,53], both of which were elevated in COMT5139 plants. Some of these phenolics such as syringic acid and caffeic acid could also have an inhibitory impact on the bacterial abundance in the rhizosphere [54,55]. Thus, the differential abundance of bacteria in the COMT5139 rhizosphere could also be attributed to the significantly lower syringic acid and higher caffeic acid in its root exudates. Conclusively, the differential availability of phenolic substrates contributes to the observed taxonomic shifts.

While both genera are known to degrade lignin-derived aromatics, our data suggest that their specific preferences and tolerance to intermediates may help explain their divergent responses to lignin pathway modification. However, further work is needed to directly test catabolic capabilities of these strains under controlled conditions using specific exudate compounds.

### Loss of COMT activity may elicit stress responses and influence plant-microbe interactions

Beyond structural and metabolic effects, the COMT5139 line also exhibited elevated levels of hydroxycinnamic acid amides (HCAAs) and proline-based dipeptides in root exudates. These metabolites are often associated with plant responses to abiotic or biotic stress, including reactive oxygen species (ROS) detoxification, osmoprotection, and immune signaling [56,57]. HCAAs, in particular, are known to accumulate following disruption of lignin biosynthesis and may act as antimicrobial agents or defense signals.

This correlation raises the possibility that lignin engineering may inadvertently trigger stress or immune responses in plants, even in the absence of pathogens [58]. While our current study did not detect changes in root microbiome composition, likely due to low sequencing depth, previous work has shown that activation of plant immunity can shape rhizosphere microbial communities [59],[60]. Therefore, the shifts we observed may result not only from metabolic changes in available carbon substrates but also from altered host regulatory pathways.

Future studies incorporating transcriptomics, immune marker assays, and microbial functional profiling will be necessary to distinguish between direct metabolic effects and host-mediated defense modulation.

### Implications for plant engineering and microbial ecology

Together, our findings highlight the broader ecological consequences of metabolic engineering in plants. While reducing lignin recalcitrance has clear benefits for generating substrates for microbial biofuel production [61], unintended changes in root exudate composition and consequently, microbiome composition could affect nutrient cycling, pathogen resistance, and symbiosis in field conditions. Compared to other engineered lines explored in this study, COMT5139 is disrupted furthest upstream in the lignin biosynthesis pathway (Figure 1). This suggests that perturbations occurring further upstream in a pathway can have more pleiotropic phenotypic effects, which in turn may have a greater likelihood of causing unintended ecological effects. This insight could be crucial for selecting targets when engineering plants for lignin modification.

This study demonstrates that changes in plant cell wall composition can influence the assembly of microbial communities in the rhizosphere in a genotype- and metabolite-specific manner. In particular, COMT mutations not only shift monolignol composition but also remodel root exudate chemical composition and microbial recruitment patterns. These effects are likely to be context-dependent and could interact with soil type, existing microbial communities, and environmental conditions. Given the positive correlation between rhizosphere microbiome and exudate metabolome, profiling root exudates in newly engineered plants might serve as a more rapid approach to predict ecological impacts.

The scope of the present study is limited to the use of a SYNCOM to examine the impacts of lignin modification on microbial assembly and composition. While SYNCOMs represents a controlled system, it may not comprehensively reflect the complex dynamics of diverse natural microbial communities, including competition for resources and the nutrient limitations, and future studies could expand these findings to test these findings in the field (although regulation around field trials of engineered plants does make this a significant undertaking). Additionally, the low read counts observed for root endophytes in this study has further restricted our understanding to the exterior root zone, which may not reflect the full suite of altered interactions.

## Conclusions

This study demonstrates that genetic engineering of lignin biosynthesis in *B. distachyon*, particularly via disruption of the COMT gene, not only alters cell wall composition but also reshapes root exudate chemical composition and the assembly of the rhizosphere microbiome. These findings establish a mechanistic link between monolignol composition, metabolite secretion, and microbial recruitment, revealing that traits selected for improved biomass processing can have broader, unintended consequences for plant-microbe interactions. As metabolic engineering becomes increasingly central to sustainable agriculture and bioenergy crop development, it will be critical to evaluate not only agronomic traits but also ecological outcomes, including impacts on root-associated microbial communities. Integrating plant synthetic biology with microbiome-aware design holds promise for optimizing both plant performance and ecosystem function.

## Acknowledgement

The authors are grateful for colleagues who provided us with the plant lines used in this study. The lignin mutant lines COMT5139, CAD4179, PMT6063, and lac5lac8 were kindly provided by Dr. Richard Sibout at National Institute of Agricultural Research (INRA), France. The transgenic line *PMT OX* was kindly provided by Dr. John Sedbrook at Illinois State University, USA. The Bd21-3 wild type is kindly provided by John Vogel, Lawrence Berkeley National Lab, USA.

The funding support was provided by Microbial Community Analysis & Functional Evaluation in Soils (m-CAFEs), an SFA led by Lawrence Berkeley National Laboratory, based upon work supported by the U.S. Department of Energy, Office of Science, Office of Biological & Environmental Research under contract number DE-AC02-05CH11231 to TRN, JCM and AM. Additional support to JCM comes from the Australian Research Council (CE230100015, IC210100047, IH230100006). Any subjective views or opinions that might be expressed in the paper do not necessarily represent the views of the U.S. Department of Energy or the United States Government. The United States Government retains and the publisher, by accepting the article for publication, acknowledges that the United States Government retains a nonexclusive, paid-up, irrevocable, worldwide license to publish or reproduce the published form of this manuscript, or allow others to do so, for United States Government purposes.

## Author Contributions

H.H.L., S.P., A.M. and J.C.M. conceptualized the study. H.H.L., S.M., N.D. and S.P. designed and conducted the experiments. P.F.A. provided the EcoFABs and performed the initial data analysis of 16s rRNA amplicon sequencing results. S.M.K. and Y.D. performed the LC-MS/MS of root exudates. S.M.K. and S.P. did the LC-MS/MS data analysis. Y.T. performed the pyro-GC/MS of root tissues. J.C.M., A. E., T.R.N. and A.M. provided the supervision. J.C.M., A.M. and T.R.N. acquired the funds for the project. S.P. drafted the initial manuscript, and SP and JCM developed the submitted version, with input from all authors, who have read, provided feedback, and approved the manuscript for publication.

